# Capsaicin and its analogues impede nocifensive response of *Caenorhabditis elegans* to noxious heat

**DOI:** 10.1101/2020.01.21.914010

**Authors:** Bruno Nkambeu, Jennifer Ben Salem, Francis Beaudry

## Abstract

Capsaicin is the most abundant pungent molecule identified in red chili peppers, and it is widely used for food flavoring, in pepper spray for self-defense devices and recently in ointments for the relief of neuropathic pain. Capsaicin and several other related vanilloid compounds are secondary plant metabolites. Capsaicin is a selective agonist of the transient receptor potential channel, vanilloid subfamily member 1 (TRPV1). After exposition to vanilloid solution, *C. elegans* wild type (N2) and mutants were placed on petri dishes divided in quadrants for heat stimulation. Thermal avoidance index was used to phenotype each tested *C. elegans* experimental groups. The data revealed for the first-time that capsaicin can impede nocifensive response of *C. elegans* to noxious heat (32°C – 35°C) following a sustained exposition. The effect was reversed 6h post capsaicin exposition. Additionally, we identified the capsaicin target, the *C. elegans* transient receptor potential channel OCR-2. Further experiments also undoubtedly revealed anti-nociceptive effect for capsaicin analogues, including ginger (*Zingiber officinale*) and turmeric (*Curcuma longa*) secondary metabolites.

## Introduction

Capsaicin is the most abundant pungent molecule identified in chili peppers, and it is widely used for food flavoring, for pepper spray in self-defense devices and recently in ointments for the relief of neuropathic pain (Szallasi and Blumberg, 1999; Yardim, 2011). Capsaicin and several other related vanilloid compounds are secondary plant metabolites (Ochoa-Alejo, 2006). Capsaicin is a selective agonist of the transient receptor potential channel, vanilloid subfamily member 1 (TRPV1) (Caterina *et al*., 1997; Cortright and Szallasi, 2004; Knotkova et al., 2008). Other vanilloids displayed similar properties [Klein *et al*., 2015; Smart et al., 2001]. Upon sustained stimulation, TRPV1 agonists elicit receptor desensitization, leading to alleviation of pain, a consequence of receptor conformational changes and subsequent decrease of the release of pro-inflammatory molecules and neurotransmitters following exposures to noxious stimuli [Jancso et al., 2008]. Interestingly, these effects have not yet been reported in *Caenorhabditis elegans* (*C. elegans*). Adult *C. elegans* consists of 959 cells, of which 302 are neurons, which make this model attractive to study nociception at physiological levels [Wittenburg and Baumeister, 1999]. *C. elegans* is especially convenient for the study of nociception as it presents a well-defined and reproducible nocifensive behavior, involving a reversal and change in direction away from the noxious stimuli [Wittenburg et Baumeister, 1999]. Bioinformatic analysis following the genome sequencing of *C. elegans*, identified genes encoding TRP ion channels with important sequence homologies to mammalian TRP channels including TRPVs [Kahn-Kirby *et al*. 2006]. Seven TRP subfamilies including TRPV analogs (e.g. OSM-9 and OCR-1-4) were identified and characterized. Furthermore, it has been established that *C. elegans* TRP channels are associated with behavioral and physiological processes, including sensory transduction of noxious heat [Glauser *et al*. 2011; Venkatachalam *et al*. 2014]. Many *C. elegans* TRP channels share similar activation and regulatory mechanisms with their mammal counterparts. Preliminary results suggest that the thermal avoidance response of *C. elegans* is increased by the application of the TRPV1 agonist capsaicin compatible with the characteristic pungent effect but did not appear to attenuate the perceived intensity of noxious heat under the experimental conditions used during these studies [Wittenburg and Baumeister, 1999; Tobin *et al*. 2002]. Though, the duration of exposure and the actual exposition levels could explain these results. Consequently, we hypothesized that a sustained stimulation with capsaicin and other vanilloid analogs will lead to receptor desensitization and will impede nocifensive response to noxious heat. The objective of this study is to characterize the capsaicin exposure–response relationship using *C. elegans* and heat avoidance behavior analysis [Nkambeu *et al*. 2019]. Selected capsaicin analogs and known TRPV1 agonists displayed in Figure 1 will be tested, including olvanil [Alsalem *et al*., 2016], curcumin [Zhu *et al*., 2014], 6-gingerol [Bhattarai *et al*., 2006], 6-shogoal [Bhattarai *et al*., 2006] as well as the TRPV1 antagonist capsazepine [Walpole *et al*., 1994]. Curcumin, 6-gingerol, 6-shogoal are plant secondary metabolites containing the vanillyl group suspected of being essential for vanilloid receptor interactions [Zhu et al., 2014; Bhattarai *et al*., 2006]. These secondary plant metabolites have fshown anti-inflammatory, analgesics, antioxidant and anti-cancer properties [Geng *et al*., 2016; Kim *et al*., 2005].

**Figure 1.**
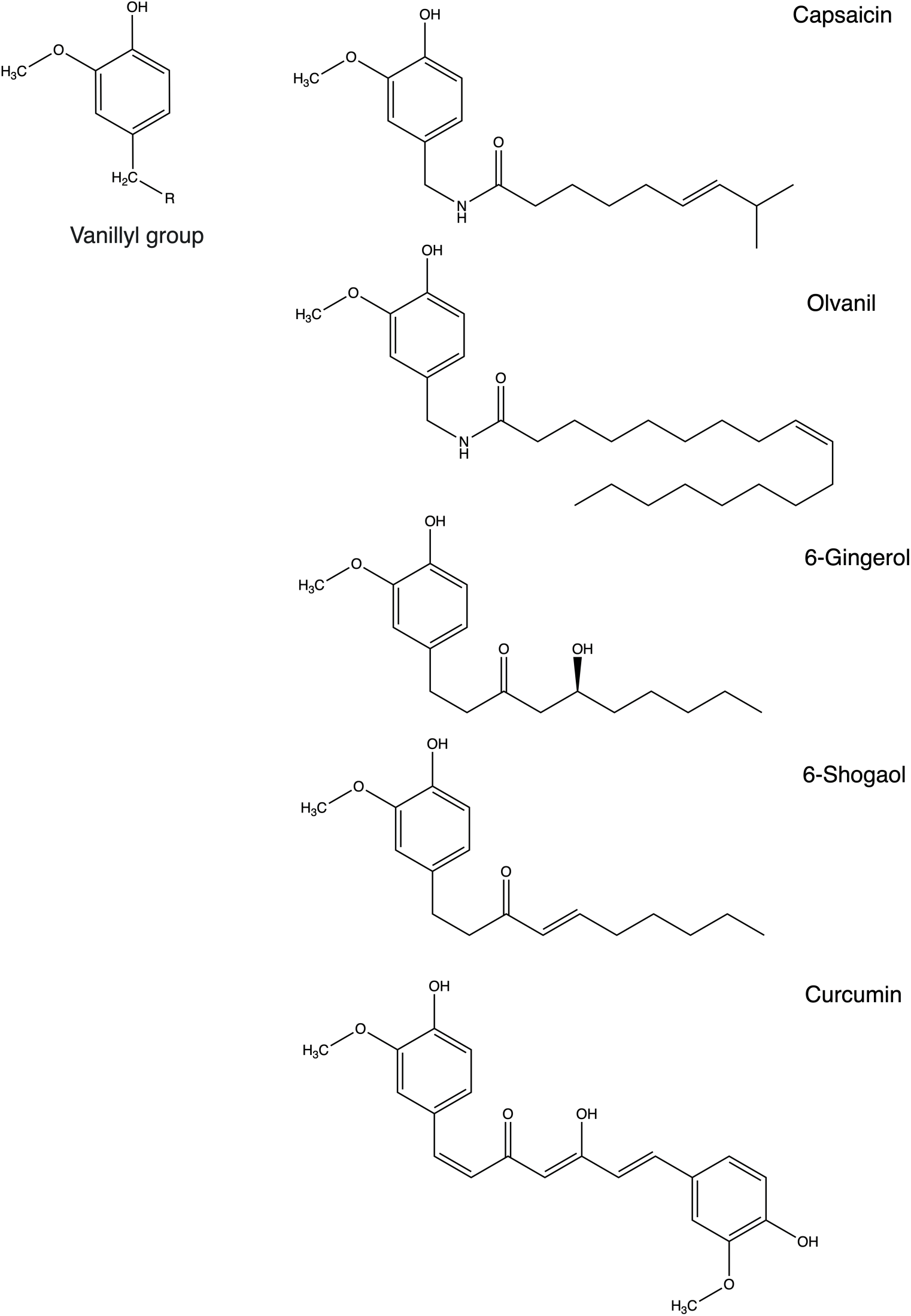
Molecular structure of capsaicin and analogues, secondary metabolites found in various peppers and spices and known TRPV1 ligands. Ligand-receptor interactions are typically associated with the pharmacophore features and vanillyl group play a central role in capsaicin and analogues specific interaction with the TRPV1.

## Materials and Methods

### Chemicals and reagents

All chemicals and reagents were obtained from Fisher Scientific (Fair Lawn, NJ, USA) or MilliporeSigma (St-Louis, MO, USA). Capsaicin, olvanil, [6]-gingerol, [6]-shogaol, curcumin and capsazepine were purchased from Toronto Research Chemicals (North York, ON, CAN).

### *C. elegans* strains

The N2 (Bristol) isolate of *C. elegans* was used as a reference strain. Mutant strains used in this work included: *ocr-1* (CX4534), *ocr-2* (JY243), *ocr-3* (RB1374), *ocr-4* (LX950) and *osm-9* (JY190). N2 (Bistrol) and other strains were obtained from the Caenorhabditis Genetics Center (CGC), University of Minnesota (Minneapolis, MN, USA). Strains were maintained and manipulated under standard conditions as described [Brenner, 1974; Margie *et al*., 2013]. Nematodes were grown and kept on Nematode Growth Medium (NGM) agar at 22°C in a Thermo Scientific Heratherm refrigerated incubator. Analyses were performed at room temperature unless otherwise noted.

### *C. elegans* pharmacological manipulations

Capsaicin was dissolved in Type 1 Ultrapure Water at a concentration of 50 µM. The solution was warmed for brief periods combined with vortexing and sonication for several minutes to completely dissolve capsaicin. Further dilution at 25 µM, 10 µM and 2 µM in Type 1 Ultrapure Water was performed by serial dilution. Olvanil, [6]-gingerol, [6]-shogaol, curcumin and capsazepine solutions were prepared using the same protocol. *C. elegans* were isolated and washed according to protocol outline by Margie *et al*. (2013). After 72 hours of feeding and growing on 92 × 16 mm petri dishes with NGM, the nematodes were off food and were exposed to capsaicin solution. An aliquot of 7 mL of capsaicin or other tested solutions was added producing a 2-3 mm solution film (solution is partly absorbed by NGM), consequently, nematodes are swimming in solution. *C. elegans* were exposed for specific times, isolated and washed thoroughly prior behavior experiments. For the residual effect (i.e. 6h latency) evaluation, after exposition to capsaicin solutions, nematodes were isolated, carefully washed and deposit on NGM free of capsaicin for 6h prior testing.

### Thermal avoidance and chemotaxis assays

The principle behind evaluating the *C. elegans* response to a stimulus (i.e. thermal or chemical) is to observe and quantify the movement evoked in response to a specific stimulus. The method proposed in this manuscript for the evaluation of thermal avoidance was modified from the four quadrants strategies previously described [Margie *et al*., 2013; Porta-de-la-Riva *et al*. 2012]. The experimental schematics are illustrated in Figure 2. Experiments were performed on 92 × 16 mm petri dishes divided into four quadrants. A middle circle delimited (i.e. 1 cm diameter) an area where *C. elegans* were not considered. The quadrants create an alternating configuration of thermal stimuli areas and control areas to prevent any bias that may appear resulting from the original position of the nematodes. Petri dishes were divided into quadrants; two stimulus areas (A and D) and two control areas (B and C). Sodium azide (i.e. 0.5M) was used in all quadrants to paralyze the nematodes. Noxious heat was generated with an electronically heated metal tip (0.8 mm diameter) producing a radial temperature gradient (e.g. 32-35°C on the NGM agar at 2 mm from the tip measured with an infrared thermometer). Nematodes were isolated and washed according to protocol outline by Margie *et al*. (2013). At this point, all nematodes tested were off food during all experimentations. The nematodes (i.e. 50 to 250 young adult worms) were placed at the center of a marked petri dish and after 30 minutes, they were counted per quadrant. Note that nematode that did not cross the inner circle were not considered. The derived Thermal avoidance Index (TI) formula is shown in Figure 2. Both TI and the animal avoidance percentage were used to phenotype each tested *C. elegans* experimental groups. The selection of quadrant temperature was based on previous experiments [Wittenburg and Baumeister, 1999].

**Figure 2.**
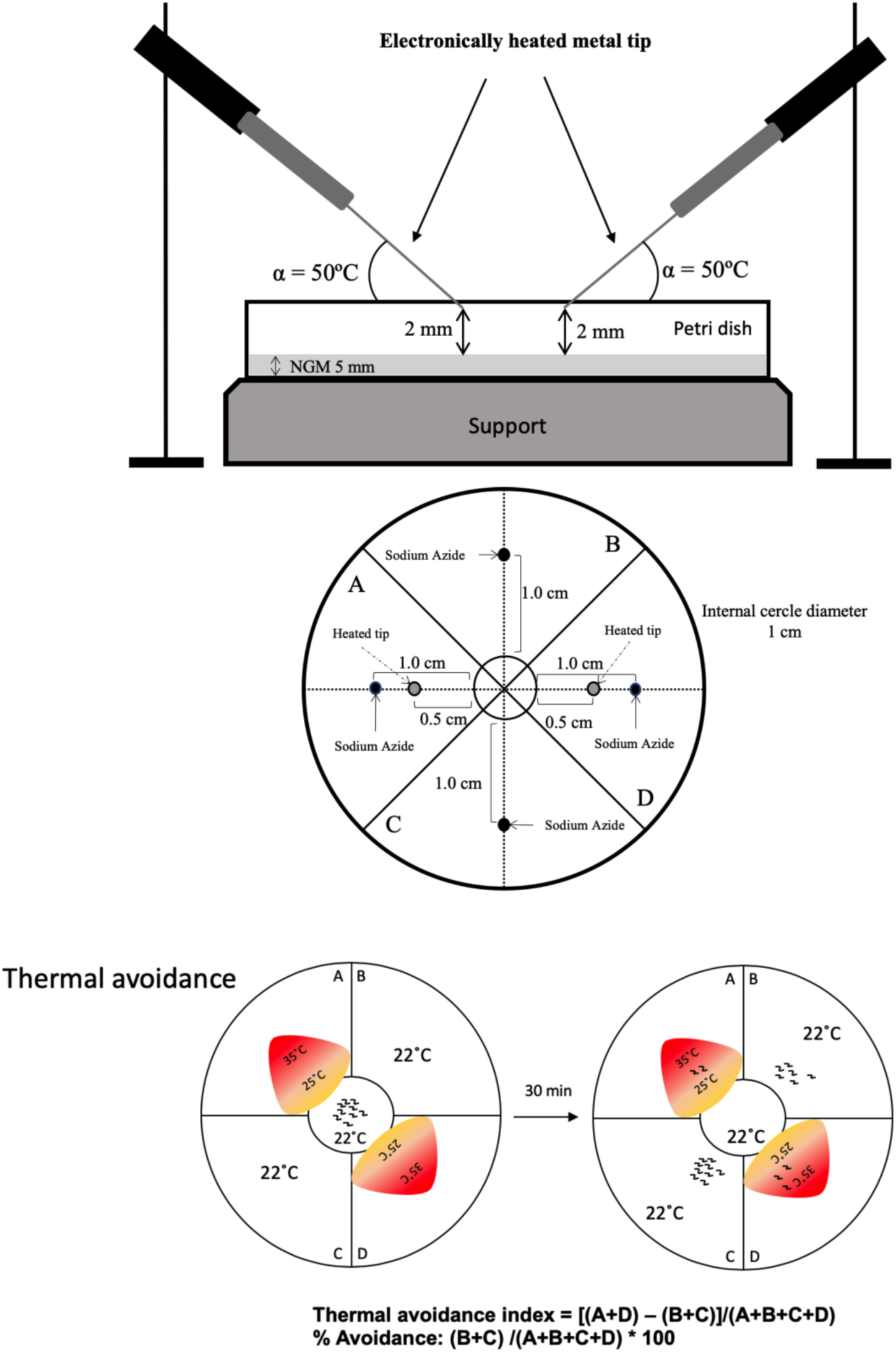
A schematic of the four quadrants assay adapted from Margie et al. (2013). For head avoidance assay, plates were divided into quadrants two test (A and D) and two controls (B and C). Sodium azide was added to all four quadrants to paralyze nematodes. *C*.*elegans* were added at the center of the plate (n=50 to 250) and after 30 minutes, animals were counted on each quadrant. Only animals outside the inner circle were scored. The calculation of thermal avoidance index was performed as described.

### Statistical analysis

Behavior data were analyzed using a one-way ANOVA followed by Dunnett multiple comparison test (e.g. WT(N2) was the control group used). Data presented in Fig 4C were analyzed using a two-tailed Student’s t-test (pairwise comparison). Significance was set a priori to p < 0.05. The statistical analyses were performed using PRISM (version 8.3).

**Figure 3.**
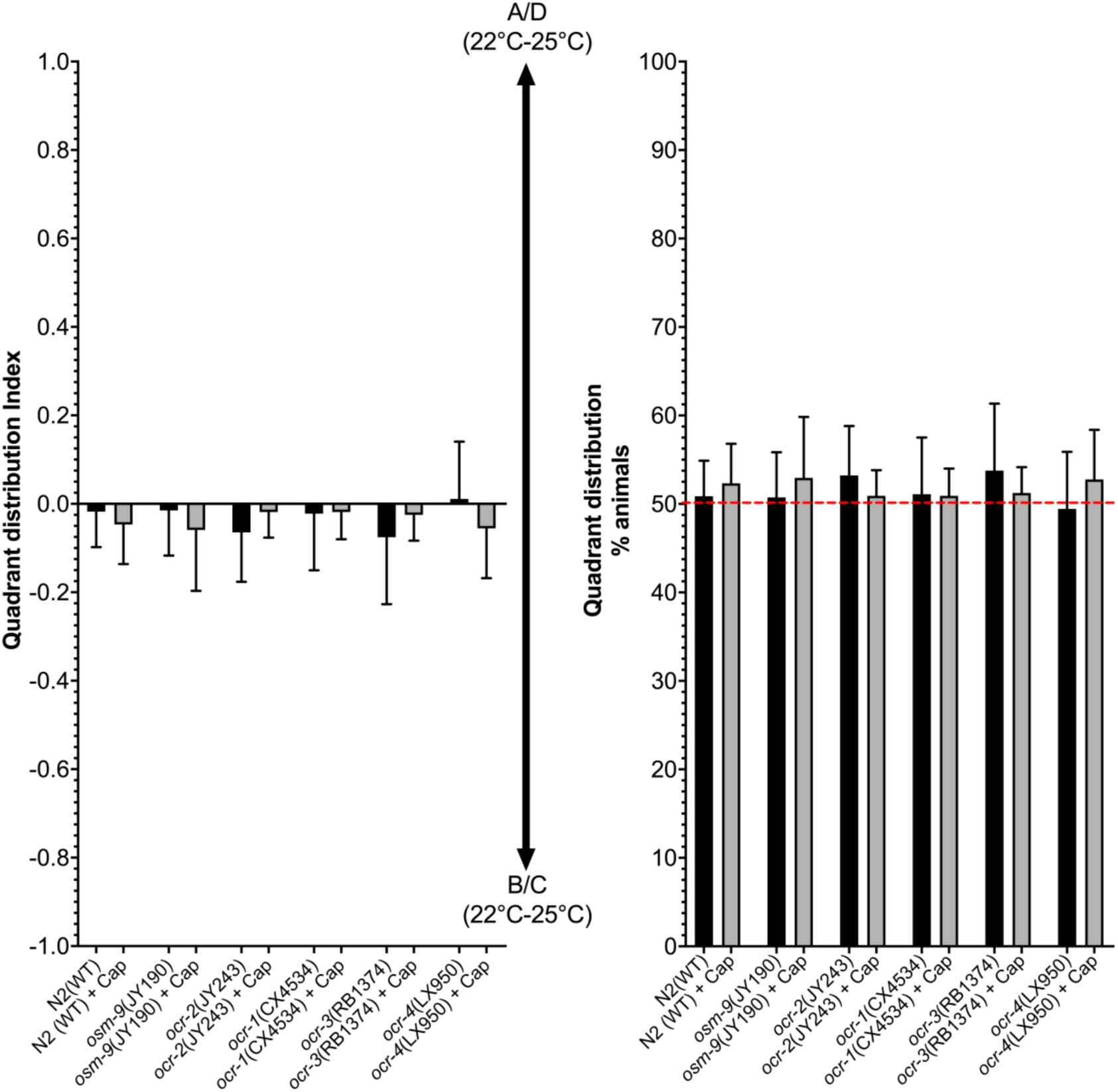
Comparison of the mobility and bias for WT (N2) and mutants *ocr-1, ocr-2, ocr-3, ocr-4* and *osm-9* nematodes in plates divided into quadrants conserved at constant temperature (22°C) and no stimulus was applied (negative control). No quadrant selection bias was observed for all *C. elegans* genotype tested in absence or presence of capsaicin at 50 µM.

**Figure 4.**
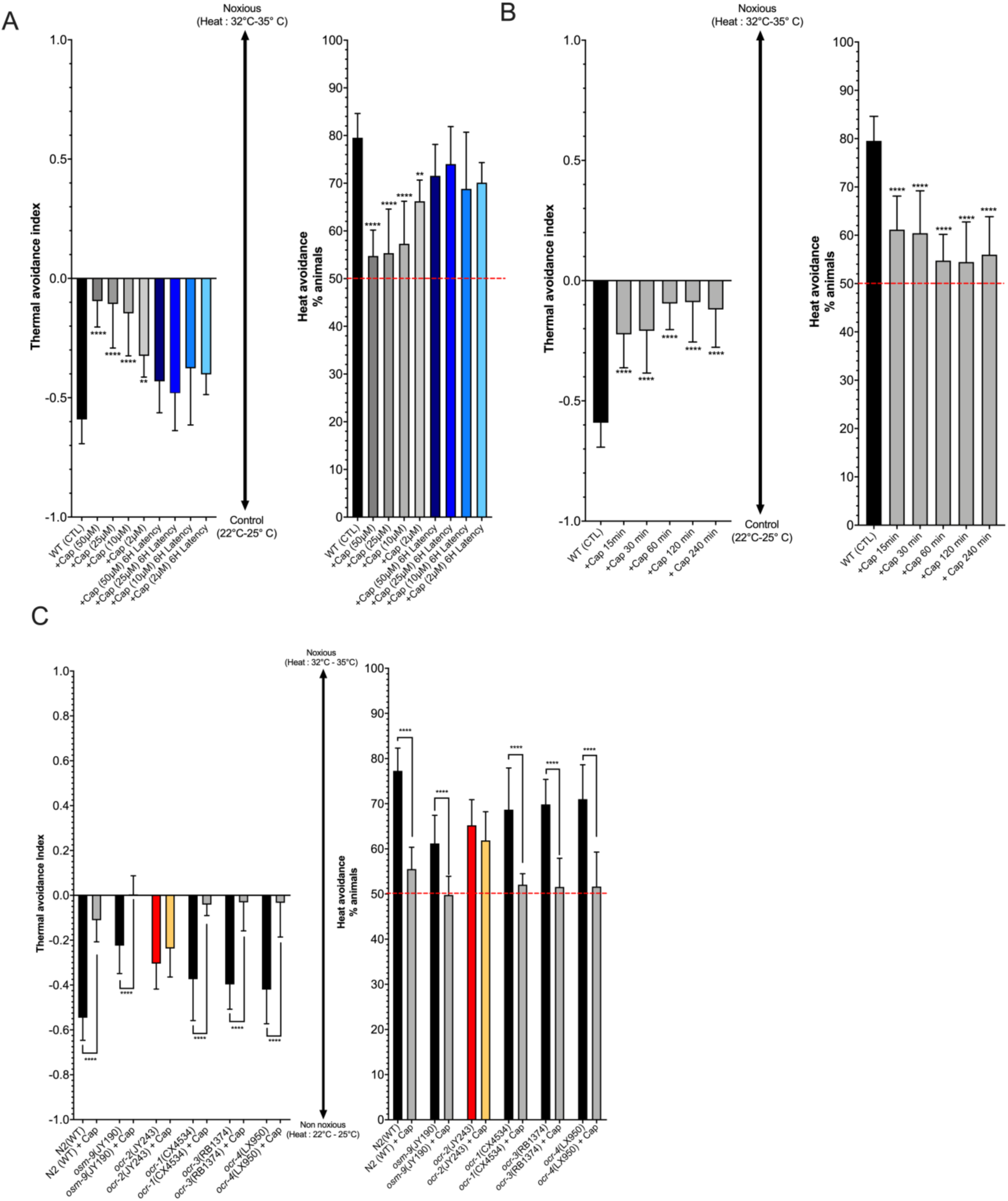
Assessment of the pharmacological effect of capsaicin on thermal avoidance in *C. elegans*. **A)** Capsaicin (Cap) dose-response assessment. Nematodes were exposed to capsaicin for 60 min prior behavior experimentations. The observed capsaicin effect is dose-dependent and noticeably impedes thermal avoidance in *C. elegans*. **** p < 0.0001, ** p < 0.01 (ANOVA - Dunnett’s Multiple Comparison). **B)** Capsaicin (Cap) time-dependent response evaluation. Nematodes were exposed to 50 µM of capsaicin in solution for various time periods prior behavior experimentations. The decrease thermal avoidance was plateaued following a 60 min exposition. **** p < 0.0001 (ANOVA - Dunnett’s Multiple Comparison). **C)** Identification of TRPV orthologs responsible for capsaicin-induced anti-nociceptive effect. Data strongly suggest that capsaicin exerts anti-nociceptive effects through the ORC-2 *C. elegans* TRPV ortholog. **** p < 0.0001 (two-tailed Student’s t-test)

## Results and discussion

Thermal avoidance assays are widely used as a model to study nociception in *C. elegans* [Glauser *et al*., 2011; Wittenburg and Baumeister, 1999]. Noxious temperatures (> 30°C) trigger a temperature avoidance response in *C. elegans* that can be quantified using a standard assay [Nkambeu *et al*. 2019; Margie *et al*., 2013; Glauser *et al*., 2011; Wittenburg and Baumeister, 1999]. Detailed studies have shown that AFD neurons are the main thermosensors in *C. elegans* [Mori and Ohshima 1995 ; Koutarou *et al*., 2004]. Besides, FLP neurons located in the head and PHC neurons in the tail act as thermal nociceptive neurons and both express heat- and capsaicin-sensitive TRPV channels, OCR-2 and OSM-9 [Liu *et al*., 2012; Glauser *et al*., 2011]. The thermal avoidance assay we have performed is described in Figure 2 and was specifically used to assess if capsaicin can impede nocifensive response to noxious heat and we tested specific *C. elegans* mutants to identify capsaicin targets. The initial experiment involved an assessment of the mobility and bias of WT (N2) and mutants *ocr-1, ocr-2, ocr-3, ocr-4* and *osm-9* nematodes in absence or in presence of capsaicin. Nematodes were placed in the center of plates divided into quadrants conserved at constant temperature (i.e. 22°C) and no heat stimulus was applied (negative control). As presented in Figure 3, there was no quadrant selection bias observed for all *C. elegans* experimental groups or genotypes tested with or without capsaicin exposition (50 µM). The nematodes were not preferably selecting any quadrant and were uniformly distributed after 30 minutes following the initial placement at the center of the marked petri dish.

Preliminary results have suggested that the thermal avoidance response of *C. elegans* is increased following *C. elegans* exposition to capsaicin [Wittenburg and Baumeister, 1999; Tobin *et al*. 2002]. As it is well described in the literature, TRPV1 agonists (e.g. capsaicin, resiniferatoxin and vanilloids) activate the TRPV1. Upon sustained stimulation, TRPV1 agonists elicit receptor desensitization, leading to alleviation of pain, which results from conformational changes, along with subsequent decrease of the release of pro-inflammatory molecules and neurotransmitters following exposures to noxious stimuli. Up to now, these effects were not reported in *C. elegans* most likely due to uncontrolled capsaicin exposition levels and time during these studies. Thus, we have exposed nematodes to capsaicin in solution and consequently had complete control of time and exposition levels. As shown in Figure 4A, data revealed a clear dose–response relationship with a significant anti-nociceptive effect following a 1h exposition to capsaicin at concentration ranging from 2 µM to 50 µM. Following capsaicin exposition, nematodes were thoroughly washed and transferred on NGM agar kept at 22°C in an incubator for 6h (i.e. residual effect/latency test) and thermal avoidance response was retested. Data suggest that after 6h post exposition, *C. elegans* thermal avoidance response returned to normal. Thus, no residual anti-nociceptive effects of capsaicin were observed after 6h. Capsaicin sustained exposition is an important factor to observe vanilloid receptor desensitization, therefore exposition time can be a determining variable. As presented in Figure 4B, capsaicin anti-nociceptive effects were observed after a 15 min exposition (p < 0.0001) at 50 µM but it is only after 60 min (p < 0.0001), we observed the highest anti-nociceptive effects at 50 µM. We can conclude that capsaicin anti-nociceptive effects are time and concentration dependent in *C. elegans*. Also, other experiments were conducted on specific *C. elegans* mutants (i.e. *ocr-1, ocr-2, ocr-3, ocr-4* and *osm-9*) to identify capsaicin target receptors. As seen in Figure 4C, capsaicin anti-nociceptive effects were quantifiable in *ocr-1, ocr-3, ocr-4* and *osm-9* mutants. However, no significant capsaicin effects (p > 0.05) were observed in *ocr-2* mutant suggesting that capsaicin target ORC-2, a transient receptor potential channel, vanilloid subfamily and a mammalian capsaicin receptor-like channel. Thermal nociceptive neurons express heat- and capsaicin-sensitive TRPV channels, OCR-2 and OSM-9 [Liu *et al*., 2012; Glauser *et al*., 2011] but our data strongly indicates that capsaicin exerts anti-nociceptive effects through a sustained stimulation with capsaicin leading to OCR-2 receptor desensitization and not the OSM-9 receptor impeding nocifensive response to noxious heat. These data sets clearly demonstrate, for the first time anti-nociceptive effects of capsaicin in *C. elegans*.

[6]-Gingerol and [6]-shogaol are major pungent components of ginger (Zingiber officinale), a plant commonly used as a spice in a variety of food preparations and beverages, and as a drug in traditional Chinese medicine (Grzanna et al., 2005). Curcumin, the active ingredient of turmeric (*Curcuma longa*), has a wide range of beneficial effects including anti-nociceptive effects in animal models and in humans [Zhu *et al*., 2014]. It has been suggested that curcumin or curcuminoid anti-nociceptive effects involved interaction with the TRPV1 receptors [Lee *et al*., 2013]. [6]-Gingerol, [6]-shogaol and curcumin are structural analogs of capsaicin, and molecular modeling studies indicate that the vanillyl moiety, as well as the long unsaturated acyl chain, have significant impacts on relative binding affinities with the TRPV1 receptor (Viswanadhan *et al*., 2007). Thus, the chemical structure similarities of [6]-gingerol, [6]-shogaol and curcumin with capsaicin suggest that it could be a good ligand of the capsaicin-sensitive TRPV channels and produce anti-nociceptive effects in *C. elegans*. Olvanil is a potent agonist of the TRPV1 and it is 10-fold more potent agonist compared to capsaicin [Alsalem et al., 2016]. Olvanil is used as a positive control in our study. Capsazepine, a well-studied TRPV1 antagonist, attenuates nocifensive responses but its efficacy strongly varies depending on the experimental model used [Walpole *et al*., 1994; Salat and Filipek, 2015]. Thermal avoidance response of *C. elegans* was tested following a 1h exposition to [6]-gingerol, [6]-shogaol and curcumin as well as olvanil and the antagonist, capsazepine all at 50 µM. The initial experiment involved an assessment of the mobility and bias of WT (N2) nematodes in absence or in presence of [6]-gingerol, [6]-shogaol and curcumin as well as olvanil and the antagonist, capsazepine all at 50 µM without noxious heat stimulation. As shown in Figure 5A, nematodes were not preferably selecting any quadrant and were uniformly distributed after 30 minutes following the initial placement at the center of the marked petri dish for all tested group. Thus, we performed the thermal avoidance test since these molecules had no impact on normal nematode behavior. The data shown in Figure 5B, revealed that all capsaicin analogues (p < 0.0001) and capsazepine (p < 0.0001) hinder nocifensive response of *C. elegans* to noxious heat. The anti-nociceptive effects observed following a 1h exposition at 50 µM is comparable to capsaicin. Anti-nociceptive effects or analgesia was observed in animal models of pain or in human for all these capsaicin analogues [Jones *et al*., 2011; Groninger and Randall, 2012; Tamburini *et al*., 2018]. Interestingly, both ginger (*Zingiber officinale*) and turmeric (*Curcuma longa*) are most likely using the phenylpropanoid pathway and generate putative type III polyketide synthase molecules responsible for pharmacological activities like anti-nociceptive effects or analgesia in higher animal species [Zhu *et al*., 2014; Gauthier *et al*., 2013].

**Figure 5.**
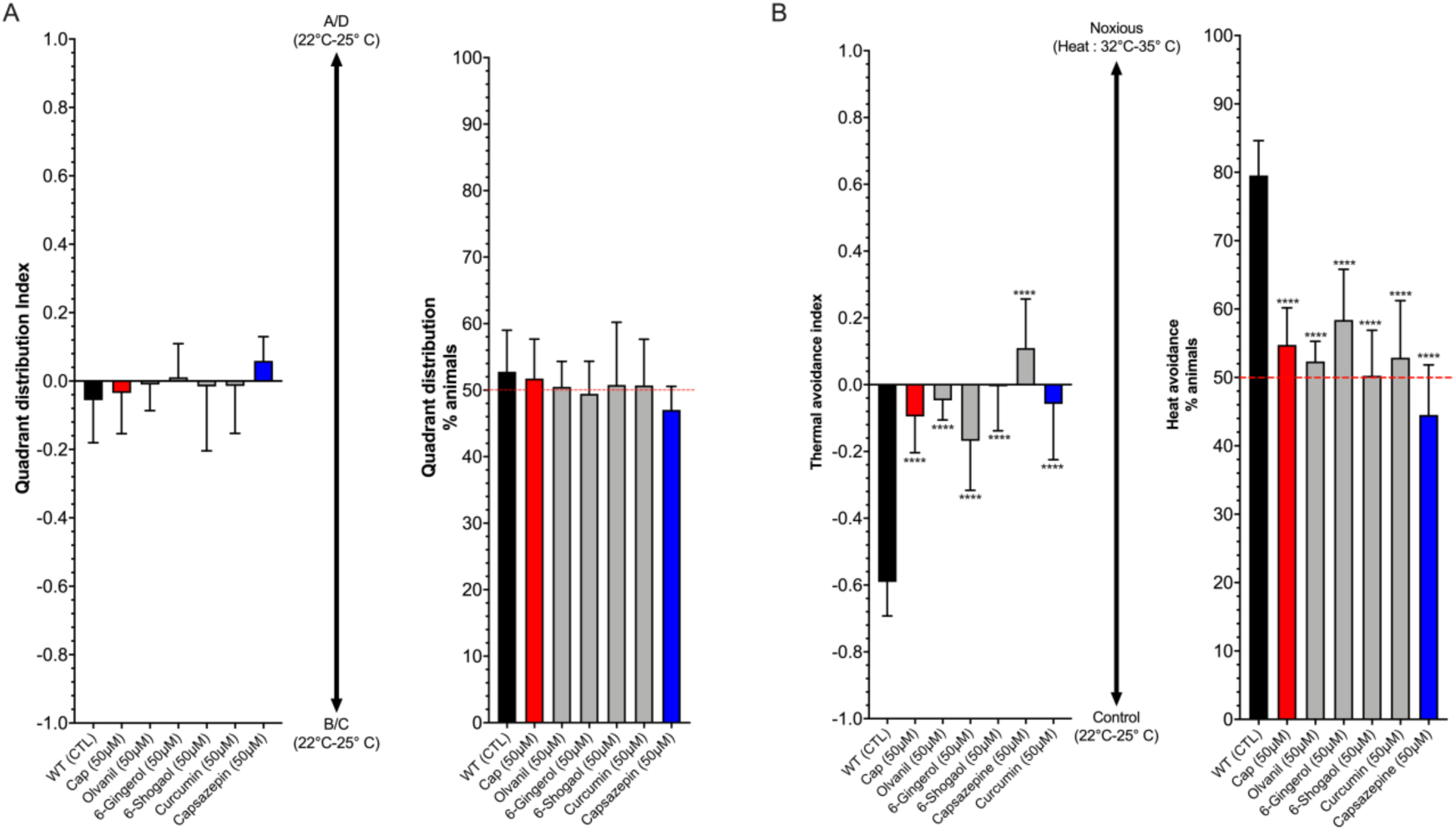
Assessment of the anti-nociceptive and desensitizing effects of vanilloid analogues of capsaicin. All tested analogues (50 µM) and capsaicin (50 µM) produced significant anti-nociceptive and desensitizing effects in *C. elegans*. Moreover, capsazepine (50 µM), a known antagonist of the TRPV1, hampered the heat avoidance behavior in *C. elegans*.

## Conclusion

This study has shown for the first-time capsaicin anti-nociceptive effect in *C. elegans* following a controlled and prolonged exposition. Additionally, we have identified the capsaicin target, OCR-2. Further experiments also undoubtedly revealled anti-nociceptive effect for capsaicin analogues, including ginger (*Zingiber officinale*) and turmeric (*Curcuma longa*) secondary metabolites. The usage of capsaicin as a clinically viable drug is limited by its unpleasant side effects, such as burning sensation, gastric irritation and stomach cramps. The rapid growing technological advancements allows the identification and isolation of molecules from specific herbal products. Using mass spectrometry, we can specifically identify and isolate secondary metabolites containing the vanillyl group, the capsaicin pharmacophore. *C. elegans* offer an opportunity for screening of large vanilloid libraries for anti-nociceptive activity to isolate rapidly effective constituents.

## Acknowledgements

This project was funded by the National Sciences and Engineering Research Council of Canada (F. Beaudry discovery grant no. RGPIN-2015-05071). Laboratory equipment was funded by the Canadian Foundation for Innovation (CFI) and the *Fonds de Recherche du Québec (FRQ)*, the Government of Quebec (F.Beaudry CFI John R. Evans Leaders grant no. 36706). A PhD scholarship was awarded to J. Ben Salem with a grant obtained from *Fondation de France*.

